# Structure of Cystathionine β-Synthase from Toxoplasma gondii, a key enzyme in its H_2_S production machinery

**DOI:** 10.1101/2021.01.15.426774

**Authors:** In memory of Jan P. Kraus, Carmen Fernández-Rodríguez, Iker Oyenarte, Carolina Conter, Irene González-Recio, Reyes Nuñez-Franco, Claudia Gil-Pitarch, Iban Quintana, Gonzalo Jiménez-Osés, Paola Dominici, Maria Luz Martinez-Chantar, Alessandra Astegno, Luis Alfonso Martínez-Cruz

## Abstract

Cystathionine β-synthase (CBS), the pivotal enzyme of the reverse transsulfuration pathway, catalyzes the pyridoxal-5’-phosphate-dependent condensation of serine with homocysteine to form cystathionine. Additionally, CBS performs alternative reactions that use homocysteine and cysteine as substrates leading to the endogenous biosynthesis of hydrogen sulfide (H_2_S), an important signal transducer in many physiological and pathological processes. *Toxoplasma gondii,* the causative agent of toxoplasmosis, encodes a functional CBS (*Tg*CBS) that contrary to human CBS, is not allosterically regulated by S-adenosylmethionine and can use both, Ser and *O*-acetylserine (OAS) as substrates. *Tg*CBS is also strongly implicated in the production of H_2_S, and thus involved in redox homeostasis of the parasite. Here, we report its crystal structure, the first CBS from a protozoan described so far. Our data reveals a basal-like fold that unexpectedly differs from the active conformations found in other organisms, but structurally similar to the pathogenic activated mutant D444N of the human enzyme.

## INTRODUCTION

Transsulfuration is the metabolic process that allows the interconversion of L-methionine (Met) and L-cysteine (Cys) through the intermediates L-homocysteine (HCys) and L-cystathionine (Cth) (1), (2). In evolutionary terms, transsulfuration consists in two routes, the “*forward*” and the “*reverse*” pathways, which operate in opposite directions and may coexist in some species, or alternatively be present as a single and irreversible pathway in other organisms. The *forward* transsulfuration enables the formation of HCys from Cys and is found in yeast (2), (3), plants (4) and enteric bacteria (5), (6). In turn, the *reverse* transsulfuration permits the synthesis of Cys from HCys, and represents the sole and irreversible source for Cys synthesis in vertebrates (2). The reverse transsulfuration involves two distinct steps. The first one is catalyzed by cystathionine ^®^-synthase (CBS) (7), which catalyzes a β-replacement reaction in which the hydroxyl group of L-serine (Ser) is replaced by HCys, yielding Cth and H2O (Fig. 1A)(Supp. Fig 1). The second step is mediated by cystathionine ©-lyase (CGL), that cleaves Cth into Cys, 〈-ketobutyrate and ammonia (8), (9). Inherited deficiency in CBS results in homocystinuria (OMIM no. 236200), a serious metabolic disorder characterized by a chronic accumulation of HCys, that is clinically manifested by connective tissue defects, mental retardation and thromboembolism (10). Besides the canonical reaction, CBS may also catalyze the endogenous synthesis of hydrogen sulfide (H_2_S) using Cys and HCys as substrates (Fig. 1A) (11). Importantly, decreased levels of H_2_S have been correlated with the development of cognitive deficits in schizophrenia (12) as well as with the degree of severity of Alzheimer disease (AD) (13). Indeed, hyperhomocysteinemia due to a deficient activity of CBS has been considered a risk factor of AD (14). Besides its relevance in physiological functions including inflammation, neuroprotection, vasodilation (15), an emerging role in protection of microbes against antibiotics (16), (17) and host-generated H_2_S in bacterial pathogenesis (18), has recently been discovered, thus opening a new window for therapies to treat microbial infections.

**Figure 1.**
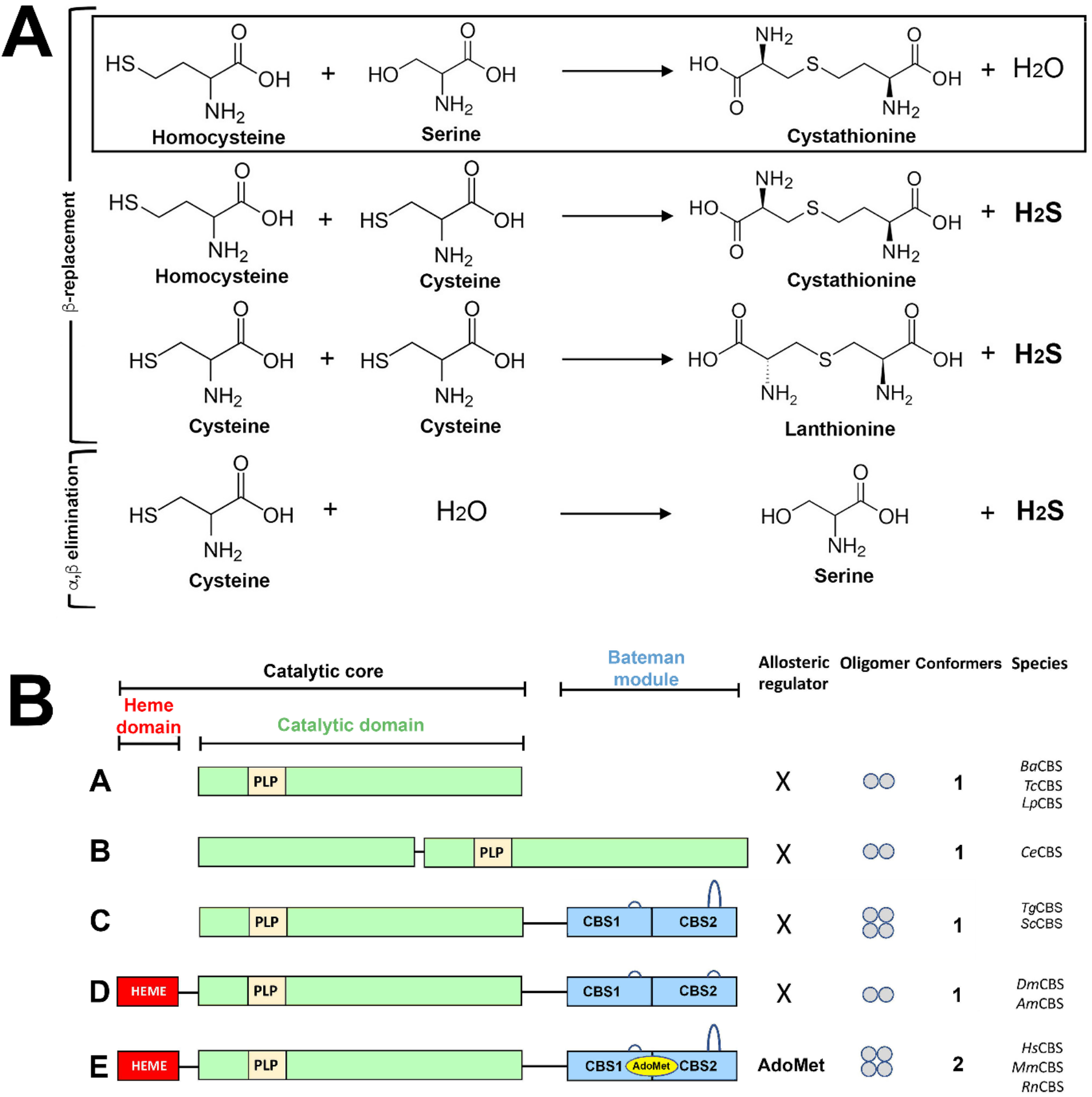
CBS reactions & architectures - Supplemental Figure 1- Supplemental Figure 2. (**A**) Reactions catalyzed by CBS enzymes. The canonical reaction is framed. (**B**) CBS architectures. Class **A**, found in *B. anthracis* (*Ba*CBS), *L. plantarum* (*Lp*CBS) and *T. cruzi* (*Tc*CBS); Class **B**, found in *C. elegans* (*Ce*CBS); Class **C,** found in *T. gondii* (*Tg*CBS) and *S. cerevisiae* (*Sc*CBS). Class **D,** found in *D. melanogaster* (*Dm*CBS) and *Apis mellifera* (*Am*CBS). All CBSs but class C, usually show a long loop at CBS2. Class **E**, found in mammals, binds AdoMet. The small circles indicate the oligomerization degree.

Structurally, CBS is a complex modular protein that has evolved adopting five different types of domain distributions (herein named as classes A to E) and two major oligomers (dimers and tetramers) (Fig. 1B) (19). The simplest architecture, or class A, is found in bacteria and consists of a catalytic domain with the fold of the type II family pyridoxal-5′-phosphate (PLP)-dependent enzymes with the cofactor covalently attached via a Schiff base bond to the ε-amino group of a conserved lysine. This type of CBS protomer self-assembles in active dimers from which reaction intermediates have been recently isolated (20), (21). A rare variant of this particular CBS, class B, is found in *Caenorhabditis elegans* (*Ce*CBS), and includes two catalytic domains in tandem, of which only one binds PLP (19), (22). The next level of complexity, class C, appears in yeast (23) and some apicomplexa (24), whose CBSs conserve the catalytic domain, and incorporate an additional C-terminal Bateman module (25), (26). The Bateman module is in turn formed by two consecutive cystathionine β-synthase motifs (CBS1, CBS2), and its actual function in these organisms remains unknown. The next CBS structural type, or class D, is found in some insects, c.a. the fruit fly (27) and the honeybee (28), and incorporates a N-terminal heme-binding domain of unknown function preceding the catalytic domain. The crystal structure of this type of CBS (27), (28), has revealed the important role played by the Bateman module in maintaining a constitutively activated dimeric assembly in these enzymes. Finally, the class E and most sophisticated version of the enzyme, is found in mammals, and consists of an allosterically regulated dimer of dimers in which each subunit is composed of three modules capable of joining three different cofactors (19) (Fig. 1). The N-terminal domain (~75 residues) binds heme and both, structural and regulatory functions have been attributed (29), (30), (31). Following the heme binding segment is the central catalytic domain, which hosts the PLP cofactor. This domain shares the fold of the type II family PLP-dependent enzymes and is the most homologous with respect to related enzymes in plants and bacteria *O*-acetylserine sulfhydrylase (OASS) and *O*-acetyl-L-serine(thiol)lyase (OASTL) (32), (33). The catalytic domain self-associates forming a compact homodimer whose structure was elucidated two decades ago (34). A long peptide linker connects the catalytic core with the third and last domain, the Bateman module, for which in mammals two main roles have been identified: i) helping to maintain the enzyme oligomerization (c.a tetramer) (35), (36), (37); ii) regulating the aperture/closure of the active site by varying its orientation with respect to the catalytic core upon binding a third cofactor, S-Adenosylmethionine, AdoMet, (37), (38). The CBS of class E, found in mammals, can adopt two different conformations. The first one, or *basal conformation,* is less active and forms basket-shaped dimers in which the Bateman module of each subunit is placed above the entrance of the catalytic cavity of its complementary counterpart, thus making it more difficult for substrates to enter the active site (38), (39). Binding of AdoMet to the Bateman module makes each protomer progress towards the so-called *activated state* (37), which is fivefold more active and associates in sea mooring bollards-shaped dimers in which the autoinhibitory effect exerted by the Bateman module is released. In 2014, we demonstrated that this *basket-to-bollard* transformation is due to a relative rotation of the two CBS motifs that weakens their interaction with the catalytic domain, and concomitantly favors their migration from above the catalytic cavity (37). This event is followed by the association of two Bateman modules from complementary subunits to form an antiparallel disk-shaped assembly known as “*CBS module*” which configures the head of the bollard and barely interacts with the catalytic domain (37). Interestingly, the enzymes of class D lack the ability of binding AdoMet and displacing the Bateman module, remaining as constitutively activated bollard-shaped dimers (27), (28). The structural knowledge of the enzyme is currently limited to the classes A, D and E described in Fig. 1.

We recently demonstrated that both CBS and CGL from *Toxoplasma gondii* (*Tg*CBS and *Tg*CGL) are functional in toxoplasma (24), (40), implying that the reverse transsulfuration pathway is operative in the parasite. In particular, *Tg*CBS can use both Ser and *O*-acetylserine (OAS) to produce Cth (24). This catalytic ability links *Tg*CBS to other enzymes such as OCBSs (41) and OASS, which also employ OAS as a substrate. Besides a role in cysteine biosynthesis, *Tg*CBS is also strongly implicated in the production of H_2_S, being highly efficient in catalyzing the β-replacement of Cys with HCys (24). Similarly to the yeast enzyme (42), *Tg*CBS belongs to the structural class C (Fig. 1), and does not respond to AdoMet stimulation (24). Herein we report the 3.2Å resolution crystal structure of an optimized protein construct (*Tg*CBSΔ466-491) that contains the full-length *Tg*CBS and only lacks amino acid residues 466–491, which belong to the long flexible loop linking the last two β-strands of the CBS2 motif. As formerly observed in *Hs*CBS (38), removal of this segment does not affect the enzyme activity. Our data, providing the first crystal structure of a full-length CBS of class C, reveal that the basal-shaped fold is not restricted to mammals and confirm that *Tg*CBS is active and forms the aminoacrylate intermediate of the β-replacement reaction either from Ser or Cys. They also explain why *Tg*CBS is not regulated by AdoMet. These results might have far-reaching consequences for the functional understanding of the molecular mechanisms involved in catalysis and regulation of other CBSs with similar domain distribution. Moreover, they pave the way for the rational design of selective drugs modulating CBS activity for use in treating toxoplasmosis.

## RESULTS

### Toxoplasma gondii CBS associates in active dimers

Careful sequence and structural alignments between *Tg*CBS and *Hs*CBS (Supp. Fig. S2) revealed an unusually extended loop (residues 463-496) between the last two β-strands of the CBS2 motif in *Tg*CBS which, as observed in humans, is presumably flexible and may impede crystal growth. Accordingly, we engineered a protein construct (*Tg*CBS Δ466-491) that lacks 25 amino acid residues of this internal loop. *Tg*CBS Δ466-491 was overexpressed in *E. coli* and purified as His-tagged protein to greater than 95% purity as judged by the presence of a single band on SDS-PAGE (Fig. 2A). Through absorbance and circular dichroism (CD) studies, we ascertained that the mutant displayed no substantial differences compared to the wild type protein (24) in either the absorbance spectroscopic features (Fig. 2B) or protein conformation (secondary structure elements) (Fig. 2C). Moreover, thermal stability studies by CD measurements at 222 nm resulted in similar denaturation profiles, indicating that the mutation did not affect the stability of the protein (Fig. 2D). The deletion of the region 466-491 did not even alter the oligomeric state of the protein since the *Tg*CBS Δ466-491 variant exists primarily as a dimer with a small amount of tetramer as observed for the wild type protein (Fig. 2E). Importantly, the kinetic parameters for canonical β-replacement reaction of Ser (or OAS) and HCys for *Tg*CBS Δ466-491 enzyme were not significantly different compared to the wild type enzyme (Supp.Table S1) (24). No response to AdoMet was observed as for the wild type protein.

**Figure 2.**
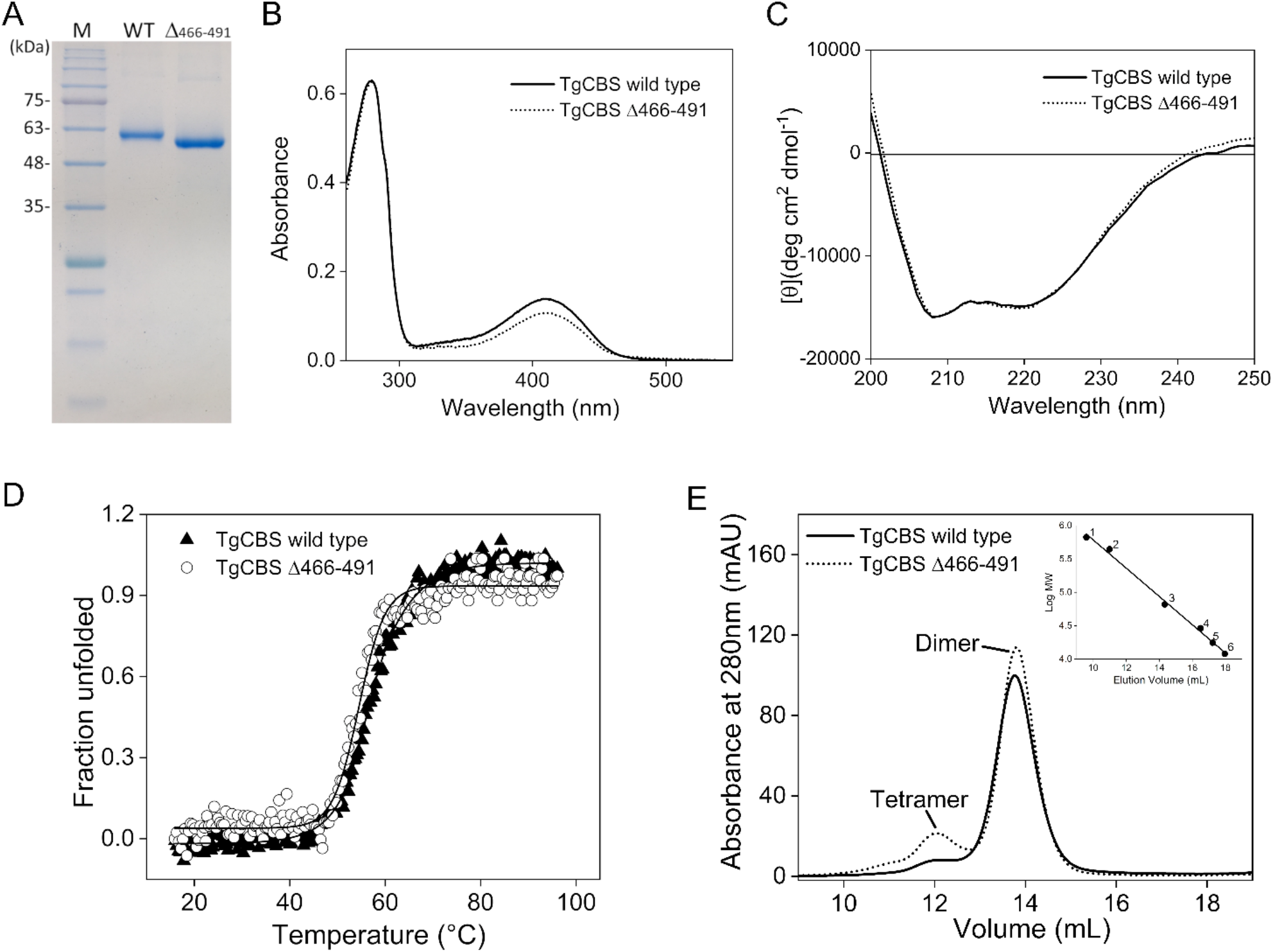
Properties of *TgCBS* Δ466-491. (**A**) 12% SDS–PAGE analysis of purified recombinant *Tg*CBS wild type and *Tg*CBS Δ466-491. Lane M, protein marker. (**B**) UV-visible absorption spectra of 15 μM purified *Tg*CBS wild type (solid line) and *Tg*CBS Δ466-491 (dotted line) recorded in 20 mM sodium phosphate buffer pH 8.5. (**C**) Far-UV CD spectra of *Tg*CBS wild type (solid line) and *Tg*CBS Δ466-491 (dotted line) at 0.2 mg/mL in 20 mM sodium phosphate buffer pH 8.5. (**D**) Thermal denaturation of *Tg*CBS wild type (solid triangles) and *Tg*CBS Δ466-491 (open circles) recorded following ellipticity signal at 222 nm at 0.2 mg/mL protein concentration in 20 mM sodium phosphate buffer pH 8.5. (**E**) Gel filtration chromatography of *Tg*CBS wild type (solid line) and *Tg*CBS Δ466-491 (dotted line) at 1 mg/mL using a Superdex 200 10/300 GL column in 20 mM sodium phosphate, 150 mM NaCl buffer pH 8.5. *Inset,* calibration curve of logarithm of the molecular weight *versus* elution volume. The standard proteins used were: 1, thyroglobulin; 2, apoferritin; 3, albumin bovine serum; 4, carbonic anhydrase; 5, myoglobin; 6, cytochrome c.

### Overall Structure of TgCBS

The *Tg*CBS Δ466-491 construct yielded crystals belonging to space groups P3_1_ and P3_1_21, and diffracting X-rays from 3.15 to 3.6 Å resolution (Supp.Table S2). In agreement with its behavior in solution, *Tg*CBS Δ466-491 forms symmetric dimers in the crystals (Fig. 3). Each protomer is composed of two independent blocks, consisting of a N-terminal catalytic domain (residues 1-323) and a C-terminal Bateman module (residues 353-514), tethered by a long linker (residues 324-352) that contains two α-helices. At first glance, the most striking characteristic is not the intrinsic folding of each of these domains, but their relative orientation, which unexpectedly reproduces the arrangement observed in the *basal conformation* of the human enzyme (Fig. 3) (38), (39). In such conformation the enzyme self-associates in dimers that adopt a basket-shaped domain-swapped symmetrical structure where the catalytic core of each protomer interacts with both, the catalytic core and the regulatory domain of the complementary monomer (Fig. 3A). Meanwhile, the regulatory domain is strategically positioned above the entrance of the catalytic crevice of the complementary monomer, thus hampering the access of substrates and retaining the enzyme in an apparent inactive state. A priori, we found these findings surprising considering the evolutionary origin of toxoplasma and the inability of *Tg*CBS to bind or to be regulated by AdoMet.

**Figure 3.**
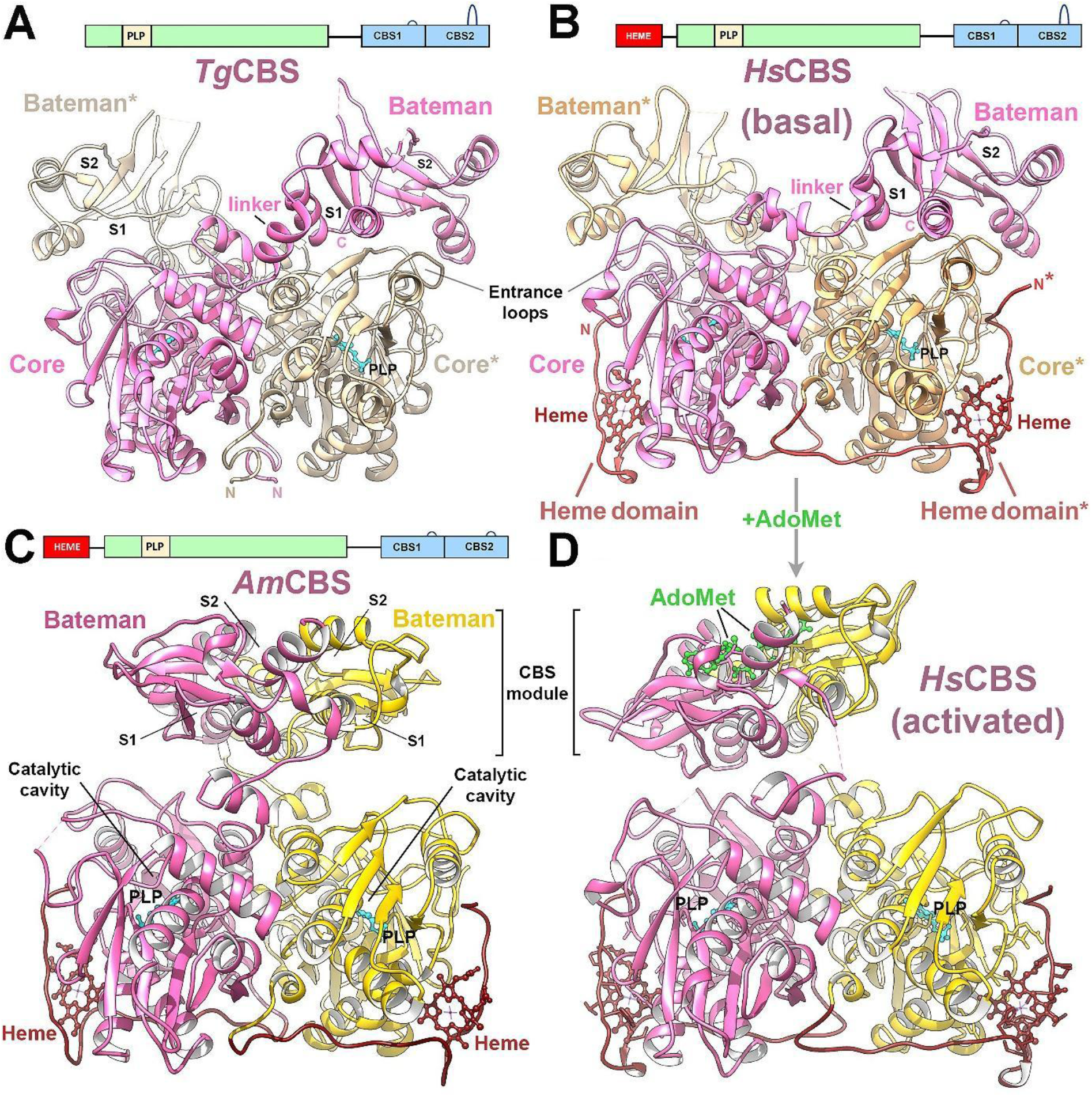
Structure of *Tg*CBS. **(A)** Structure of the basket-shaped *Tg*CBS dimer. Each subunit is colored in *yellow* and *pink*, respectively. (**B**) Basket-shaped *Hs*CBS dimer (*Basal state*) (PDB ID 4L0D). The heme-binding domain is in *red*. (**C**) Constitutively activated *Sea bollard-shaped* dimer from *Apis mellifera* CBS *(Am*CBS). (**D**) AdoMet-bound *sea bollard-*dimer of *Hs*CBS (*activated state*). The Bateman modules associate forming a disk-like *CBS module*. Heme, PLP and AdoMet are in *red*, *cyan* and *green* sticks, respectively.

Structurally, the catalytic domain of *Tg*CBS presents the overall fold of the β-family of the PLP-dependent enzymes and maintains its secondary elements basically unaltered with respect to the equivalent region in the human (Figs. 3B, D) (29), (34), (38) fly (27) and honeybee enzymes (Fig. 3C), (28). The catalytic domain is composed of fourteen α-helices and two β-sheets consisting of four (β3–β6) and six (β1–β2, and β7–β10) strands, respectively (Fig. 3A). An additional β-strand precedes the last α-helix (α14) of the domain. A main difference with the human and insect proteins, but similar to the yeast enzyme (19), is the absence of a heme-binding domain (Figs. 1, 3) (24). Two distinct blocks are clearly distinguishable in the catalytic domain (Fig. 4A). The larger one contains two segments of residues distanced in the peptide chain (residues 10-55 and 163-322) and ranges the so-called *static subdomain*. This block behaves as a rigid body during the catalysis in other CBSs and relative enzymes, and represents the region that most interacts with other protein domains (28). In *Tg*CBS, it contributes significantly to maintaining the integrity of the dimer by providing hydrophobic residues (c.a L18, V27, M29, I89, L93, V97, I117, C120, L121, I277, L282, M320, I324) at the interface between complementary subunits. The overall architecture of the static subdomain is mostly conserved when compared with the wild type human enzyme. A least square superposition of the Cα-positions of this region between *Tg*CBS and *Hs*CBS yields a root mean square deviation of only 0.48 Å and 0.58 Å at the monomer and dimer level, respectively. Importantly, this static block configures half the entrance to the catalytic site (Fig. 4A) (27),(43) and contains four out of the five consensus sequences that participate in the interaction with the different CBS substrates (blocks B2, B3, B4 and B5 in Fig. 4A) (41). The main differences with the human protein are in the region comprising residues 232-238 of block B3 (Fig. 4A), which is more directed towards the catalytic cleft in *Tg*CBS, due to a somewhat longer helix α10 (residues 228-234). The second subdomain of the catalytic core is the *mobile subdomain* (28) which is smaller in size (residues 75-157) and is intercalated in the static block, to which is linked by two loops that connect helix α4 with strand β4, and strand β7 with helix α8, respectively (Fig. 4A). The mobile subdomain provides the second half of the walls of the catalytic cavity, and acts as a cap that limits the access of substrates into the narrow channel where the PLP molecule is covalently attached via a Schiff base bond to the ε-amino group of lysine (K56) (44). Several H-bonds anchor the PLP molecule to the protein matrix and orient the cofactor appropriately within the cavity. Among them are those formed between the nitrogen of the pyridine and the Oγ of residue S287, and between the 3’-hydroxyl group of PLP and the N_δ2_ of residue N86 (Fig.4A). Importantly, N86 is coplanar with the pyridine ring of PLP to allow the appropriate ring tilt upon transaldimination (34). In the opposite side of PLP, the phosphate moiety interacts with the so-called *phosphate binding loop* (block B2 in Fig. 4A, residues 193-197), which is located between strand β8 and helix α9. This loop includes two important threonine residues (T194 and T197) as well as three glycines that form a network of H-bond interactions that ensure the correct orientation of PLP. Mutation of the equivalent threonines in *Hs*CBS is known to cause a loss of CBS activity, and impairs carbonylation of the heme moiety (31). Importantly, in *Tg*CBS the mobile subdomain collapses towards the catalytic cleft pushed by the Bateman module, which is placed just above (Fig. 4A). A similar arrangement is observed in the basal state of the human enzyme (38),(39) (Fig. 3B), where the loops L145–148, L171–174, and L191–202, defining the entrance of the catalytic site, are sandwiched between the core and the regulatory module, thus impairing free access of substrates into the PLP site. Finally, the C-terminal domain of *Tg*CBS comprises a Bateman module (~20 kDa) made up of two CBS motifs (CBS1, residues 354-425; CBS2, residues 426-514) that are connected to each other by a linker of eight amino acid residues (421-428). Both, CBS1 and CBS2 show the canonical βαββα sequence of secondary elements usually found in these type of motif (25), and contact each other via their three-stranded β-sheets. As observed in humans (38), (39), the Bateman module of *Tg*CBS presents two symmetrical cavities (S1, S2) (Fig. 4B) of which only S2 is exposed to solvent. Site S2 conserves some of the residues that would be necessary to stabilize one AdoMet molecule inside (c.a. T501 and D504 that would interact with the ribose ring of the nucleotide, or I503 and V499 that would help in accommodating the methionyl group of AdoMet), but misses other key features to host AdoMet. Among them is a longer peptide segment comprising residues 361-368 (417-423 in *Hs*CBS, Fig. 4D), that stands out preceding the strand β11b, and displaces residue T364 away from the center of the S2 cavity, thus preventing its potential interaction with the ribose ring of AdoMet. The extension of this segment distorts the hydrophobic pocket otherwise required to accommodate and orient the adenine ring of AdoMet inside the cavity (37). Additionally, a negatively charged glutamate at position 390 invades the position that the AdoMet adenine ring would potentially occupy (Figs. 4C, D). All these features disable *Tg*CBS from binding AdoMet.

**Figure 4.**
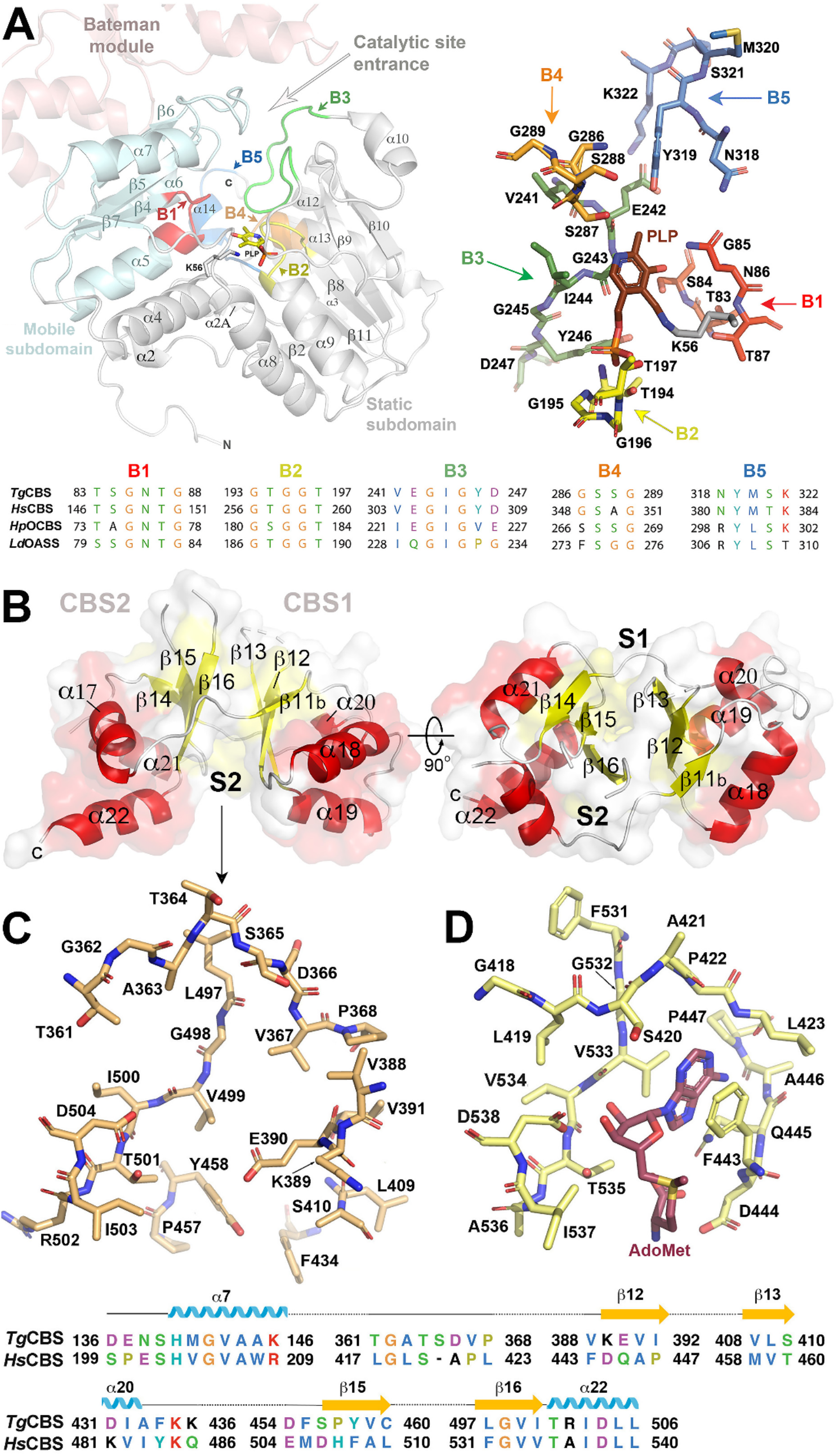
*Tg*CBS domains. (**A**, *left*) The catalytic domain of *Tg*CBS includes the *mobile* (*cyan*) and the *static* (*grey*) subdomains. The Bateman module is in *light red*. (**A**, *right*) Residues configuring the catalytic cavity. PLP (*brown*) is covalently bound to K56. (**A**, *bottom*) Sequence alignment of blocks configuring the active site. Block B1 (*asparagine loop, red*), stabilizes the aminoacrylate intermediate and contains a conserved serine (S84) in all CBSs and OASS enzymes. Block B2 (*phosphate loop, yellow*) anchors the phosphate moiety of PLP. Block B3 (*green)* is thought to interact with the second substrate, HCys, and participates in the nucleophilic attack on aminoacrylate, in the regeneration of the internal aldimine, and in the closure of the active site (41). Block B4 (*orange*) stabilizes the pyridine ring of PLP. Block B5 (*blue*) interacts with the lip of the active site. (**B**) Bateman module of *Tg*CBS. S1 and S2, are the main cavities. (**C**) Residues within cavity S2 in *Tg*CBS; (**D**) Residues in site S2 of *Hs*CBS. AdoMet is in *wine* sticks. *Bottom*: Sequence alignment of residues at cavity S2 in *Tg*CBS and *Hs*CBS.

### TgCBS exists in a unique basal-like conformation

A careful comparative analysis of the available three-dimensional data has led us to conclude that the basal-type folding of *Tg*CBS represents its sole conformational state. We base our conclusion on three key structural features. The first one is the inability of *Tg*CBS to bind AdoMet at the site S2 of the Bateman module (Fig. 4C), and thus to suffer an AdoMet-induced relative rotation of its two CBS motifs. In allosterically regulated CBS enzymes as *Hs*CBS, such rotation weakens the interactions that maintain the Bateman module anchored to the catalytic core just above the entrance of the catalytic site, and this makes the enzyme progress towards the activated state (37). It is well established that the migration of the Bateman module allows the free access of substrates into the PLP cleft (37). Nevertheless, such displacement of the Bateman module can only occur if its contacts with the catalytic core are not too tight. If this were the case, both domains would remain permanently anchored to each other (as it happens in *Tg*CBS). The second relevant feature refers to the peptide linker connecting the Bateman module with the catalytic core, which needs to be sufficiently long and flexible as to facilitate the migration of the regulatory domain to its final location in the activated state. Our structures show that, although the length of the *Tg*CBS linker is sufficient, the interactions that it maintains with the Bateman module and with the core are stronger than in *Hs*CBS (Figs. 5A, B). These contacts significantly enhance the stability of the basal conformation vs other possible arrangements, and occur thanks to a specific turn of the interdomain linker in the parasite enzyme, around residues 342-344, that allows positioning helix α16 closer to helices α6, α21 and α22 and to strands β5 and β6, than in the human protein (Fig. 5B). This peculiar location of helix α16 in *Tg*CBS results in the formation of a multiple network of salt bridges participated by residues K341, E344 and R345 (α16), K119 (α6), D454 (α21), R502 (α22), R101 (β5), and E124 (β6). These attractive electrostatic interactions complement the effect of the hydrophobic pocket formed by residues F349 from helix α16 and residues L505 and L509 from helix α22, and keep the interdomain linker of *Tg*CBS firmly attached to both the catalytic core and the Bateman module, making the displacement of the latter highly unlikely.

**Figure 5 - Supplemental Figure 3.**
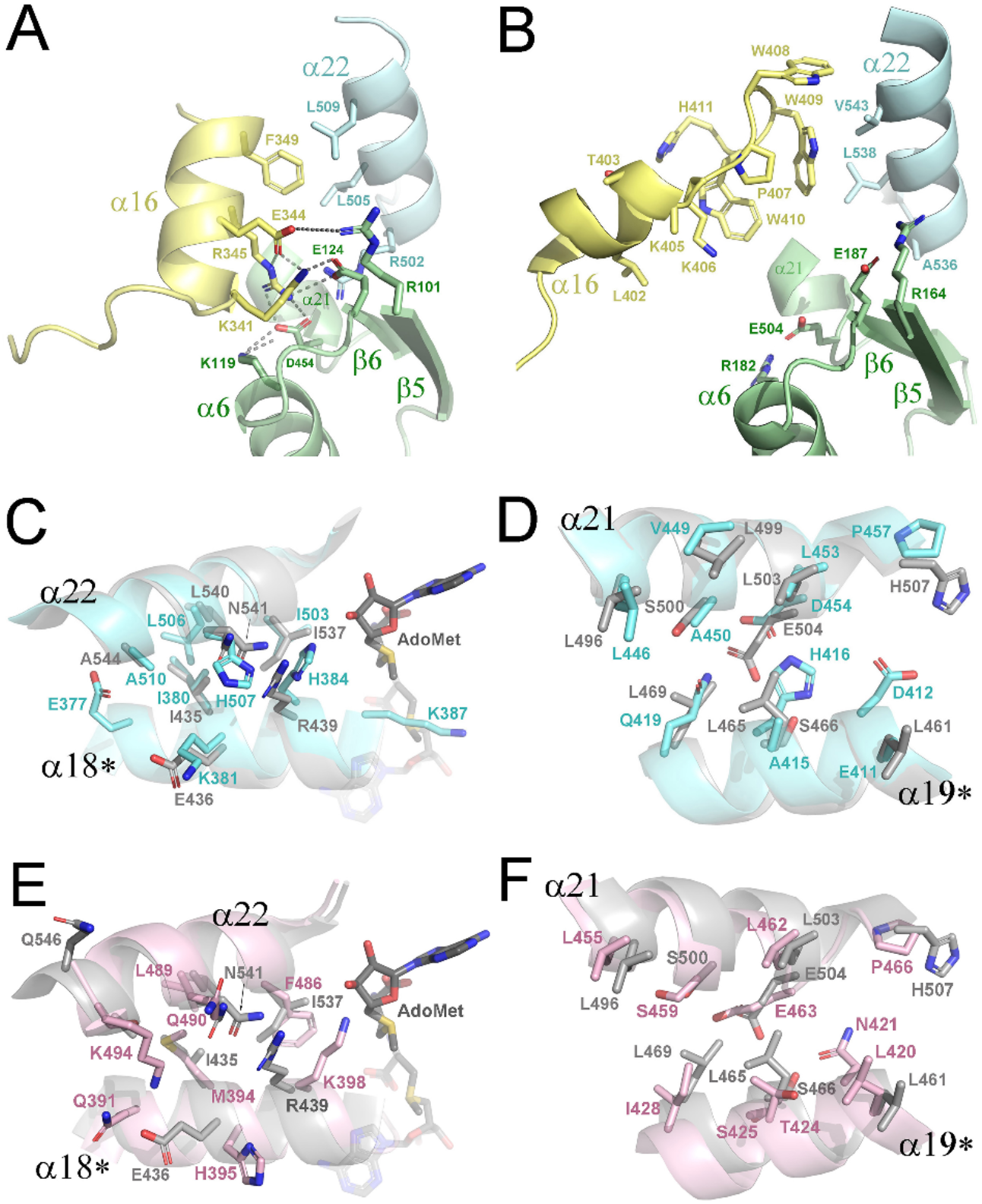
(A) Interdomain interactions in *Tg*CBS, *Hs*CBS and *Am*CBS. Interactions between the interdomain linker (*yellow*), the catalytic core (*green*) and the Bateman module (*cyan*), in *Tg*CBS (**A**) and in *Hs*CBS (**B**). Interface residues in a potential CBS module of *Tg*CBS (*cyan*) (**C**, **D**), and *Am*CBS (*pink*) (**E**, **F**) compared to that found in *Hs*CBS (*grey*) (37).

Finally, the third requirement in an allosterically regulated CBS enzyme to reach the activated state consists in being able to couple its Bateman module with the equivalent region from the complementary subunit to form a flat disk-shaped structure, known as “*CBS module*”. Once formed, this assembly locates far from the entrance of the catalytic cavity, thus allowing free access of all substrates (Supp. Fig. S3) (37). The formation of the CBS module necessarily requires the previous rotation of the CBS motifs mentioned above, which only occurs upon binding of one AdoMet molecule to the S2 site (Fig. 3) (37). The formation of the CBS module also involves establishing new interactions between the interfacial helices of the CBS domains (α18, α19 from CBS1; α21, α22 from CBS2) (Fig. 5C-F), (Supp. Fig. S3). It is worth mentioning that in all known CBS enzymes, the CBS module is antiparallel (the two Bateman modules are oriented in a head-to-tail manner) (27), (28), (37). In the said arrangement, the CBS1 of the first subunit interacts with the CBS2 motif of the complementary monomer and vice versa (Supp.Fig. S3). Obviously, if the residues at the interfacial positions do not favor the corresponding interactions, the CBS module cannot be assembled. Said this, we found that *Tg*CBS does not fulfill the third condition for activation. The independent structural superimposition of the CBS1 and CBS2 domains of *Tg*CBS and *Hs*CBS or *Tg*CBS and *Am*CBS are good (rmsd 0.902 /0.816 and 1.147/0.719, respectively), but significantly worsens when the structural alignment is performed comparing the entire Bateman module of *Tg*CBS with the corresponding region from the AdoMet-bound activated form of the human enzyme (rmsd=1.37) or to the constitutively activated *Am*CBS (rmsd=4.55). In its native conformation, the Bateman module of *Tg*CBS is not compatible with the assembly of a CBS module, as its interfacial helices would not face those of its complementary counterpart (Supp. Fig. S3). Self-assembly of Bateman modules to form a CBS module of *Tg*CBS would cause clashes between some residues (c.a K387) and a bound adenosine derivative (Fig. 5C). In such hypothetical CBS module, some favorable hydrophobic interactions could be potentially established between residues of the interfacial helices α18, α19, α21 and α22 of complementary CBS1 and CBS2 motifs (c.a L446, V449, L453 from α21; Q419, A415 from helix α19), but would also be impaired by steric clashes and/or repulsive forces between bulky or charged residues of the complementary subunits (c.a H507/H384) (Fig. 5C). Put together, all these findings strongly suggest that the basal-like fold of *Tg*CBS represents its unique conformation, and that the enzyme does not progress to a second activated form as found in mammals (37) and insects (27), (28).

### The basal-like fold of TgCBS is catalytically active

At first sight, the basal-like architecture of the crystallized *Tg*CBS and its closed active site appear to reflect an inactive form. However, detailed 3D-alignments of *Tg*CBS with *Hs*CBS revealed a slight displacement of the complementary Bateman modules towards the central cavity existing between subunits (Fig. 6). This shift also affects helices α18 and α22, and occurs without significant changes in the catalytic core (Fig. 6A). Interestingly, a similar shift was formerly observed in the pathogenic D444N mutant of *Hs*CBS (Fig. 6B) (38), which shows an approximately two-fold increase in basal activity and impaired response to AdoMet stimulation as compared with the wild type enzyme (45). Taking into account that crystallographic data offer an average spatial-time snapshot of the protein structure captured in a local energy minima under specific buffered conditions, we found reasonable to postulate that, as formerly observed in the D444N variant (38), the mentioned displacement of the Bateman module provides a greater dynamic freedom to the mobile subdomain of *Tg*CBS that facilitates the access of the substrates to the interior of its catalytic site. To confirm this hypothesis, we evaluated the conformational behavior of both, the D444N human variant and *Tg*CBS through microsecond molecular dynamics(μs-MD) simulations (Supp. Movies). To identify the most relevant conformational transitions occurring in each system, the dimensionality reduction technique of principal component analysis (PCA) was applied. Rotation and opening-closing motions of the Bateman modules were found to dominate the first two PCA’s for all the systems studied. To more precisely characterize the dynamic opening of the Bateman module, the angle between the α-carbons of residues K441, S387 of the first subunit, and K481 of the second subunit of the human enzyme was measured along the MD trajectories. The equivalent angle between residues T386, S325 and D431 of *Tg*CBS was analyzed. As hypothesized, we found that mutant D444N (Fig. 6D) and *Tg*CBS (Fig. 6E) exhibited a significantly wider opening than the wild type human enzyme (Fig. 6C). More interestingly, such opening and rotating movements of the Bateman module significantly influenced the structural elements defining the entrance of the active site, and led to the simultaneous opening of the cavity in both D444N and *Tg*CBS (Fig. 6). These effects were significantly weaker in wild-type *Hs*CBS, which explains its lower basal activity. To further explore the impact of conformational dynamics on substrate accessibility to PLP, the appearance of transient access tunnels along the MD trajectories was also analyzed. We found that mutant D444N displays a wider channel cluster than the wild-type protein, and that *Tg*CBS behaves similarly to the D444N mutant (Figs. 6C-E), (Supp. Movies). Intriguingly, the channel clusters were not symmetrically distributed between the two complementary subunits in any of the analyzed dimers, thus suggesting synergistic behavior between the two active sites. To confirm that the crystallized enzyme is indeed catalytically active, we also crystallized *Tg*CBS with each of two substrates, Ser and Cys. In agreement with the MD studies, the Polder omit maps showed additional electron density near the PLP molecule, consistent with the synthesis of the aminoacrylate intermediate. Of note, formation of the external aldimine in *Tg*CBS did not result in major distortions in the overall protein fold, and only caused subtle reorientations in the side chain of lysine K56, otherwise associated with PLP in the apoenzyme, and threonine T87, that rotates slightly to avoid clashes with the carboxylate moiety of aminoacrylate (Fig. 7). These small changes contrast with the behavior of other CBSs, such as the fly enzyme, where three loops (residues 119-122; 144-148 and 165-176) are displaced towards the PLP molecule upon addition of substrates. Formation of the aminoacrylate intermediate in the *Tg*CBS crystals is consistent with the behavior of the protein *in vitro*, where the addition of Ser or OAS results in the disappearance of the 412 nm-peak, representing the internal aldimine in the ketoenamine form, and the appearance of a major band centered at 460 nm in both absorbance and CD spectra, that is attributed indirectly to the aminoacrylate reaction intermediate (Fig. 7) (24).

**Figure 6.**
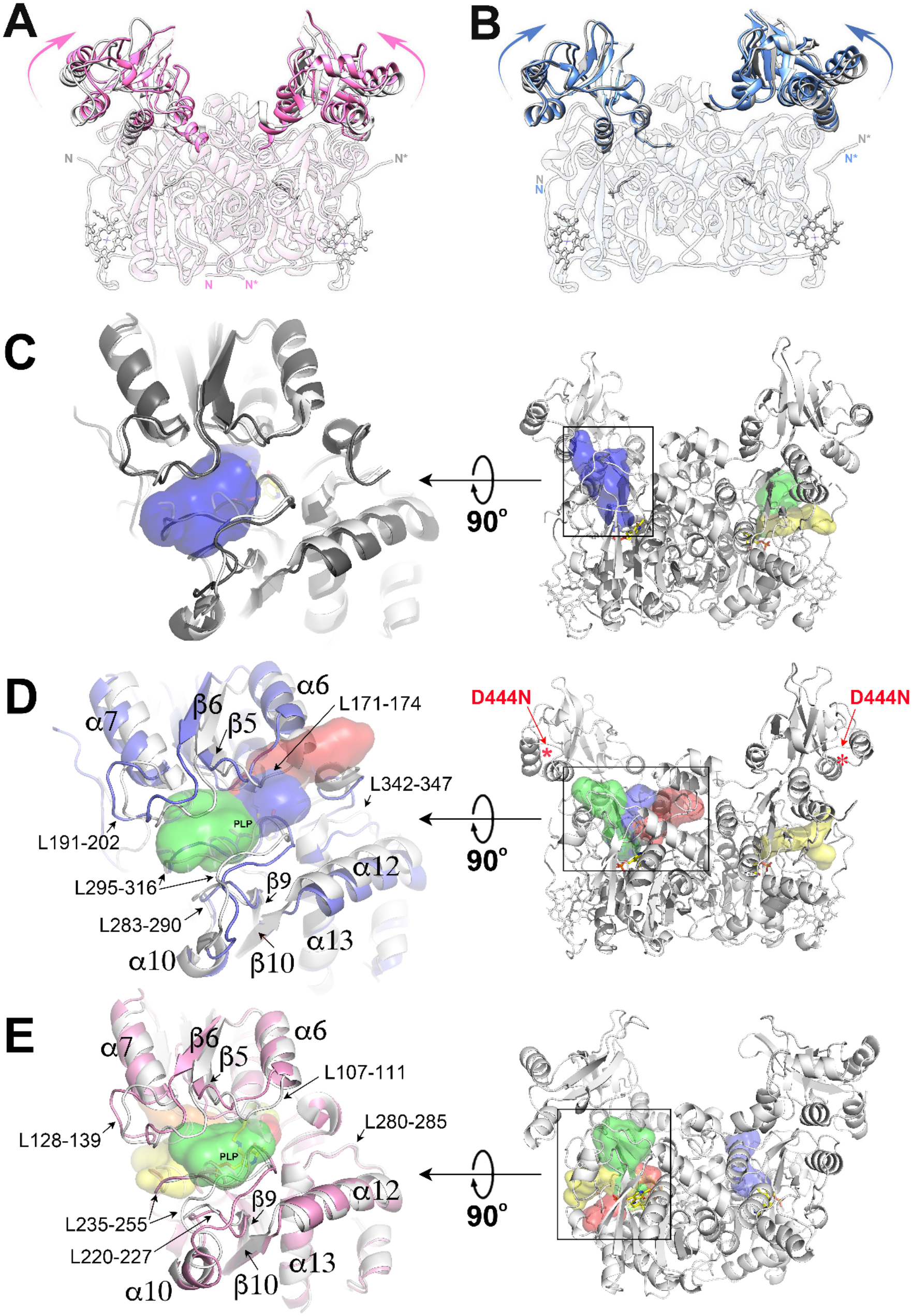
MD simulations on *Hs*CBS and *Tg*CBS. Superimposition of (**A**) *Tg*CBS (*pink*) on basal *Hs*CBS (*grey*) and (**B**) of mutant D444N_*Hs*CBS (*blue*) on basal *Hs*CBS (*grey*), respectively. The Bateman modules are in opaque ribbons. Heme and PLP are in sticks. (**C-E**) Structural overlap of the loops defining the entrance of the catalytic site at the narrower (*light grey*) and the wider amplitude points of the MD simulation in (**C**, *left*) basal wild-type *Hs*CBS, (**D**, *left*) mutant D444N_*Hs*CBS and (**E**, *left*) *Tg*CBS. *Hs*CBS, D444N_*Hs*CBS and *Tg*CBS at the wider amplitude are colored in *black*, *marine* and *pink*, respectively. The clusters of tunnels formed along the MD simulation at the corresponding catalytic sites are shown on the right.

**Figure 7.**
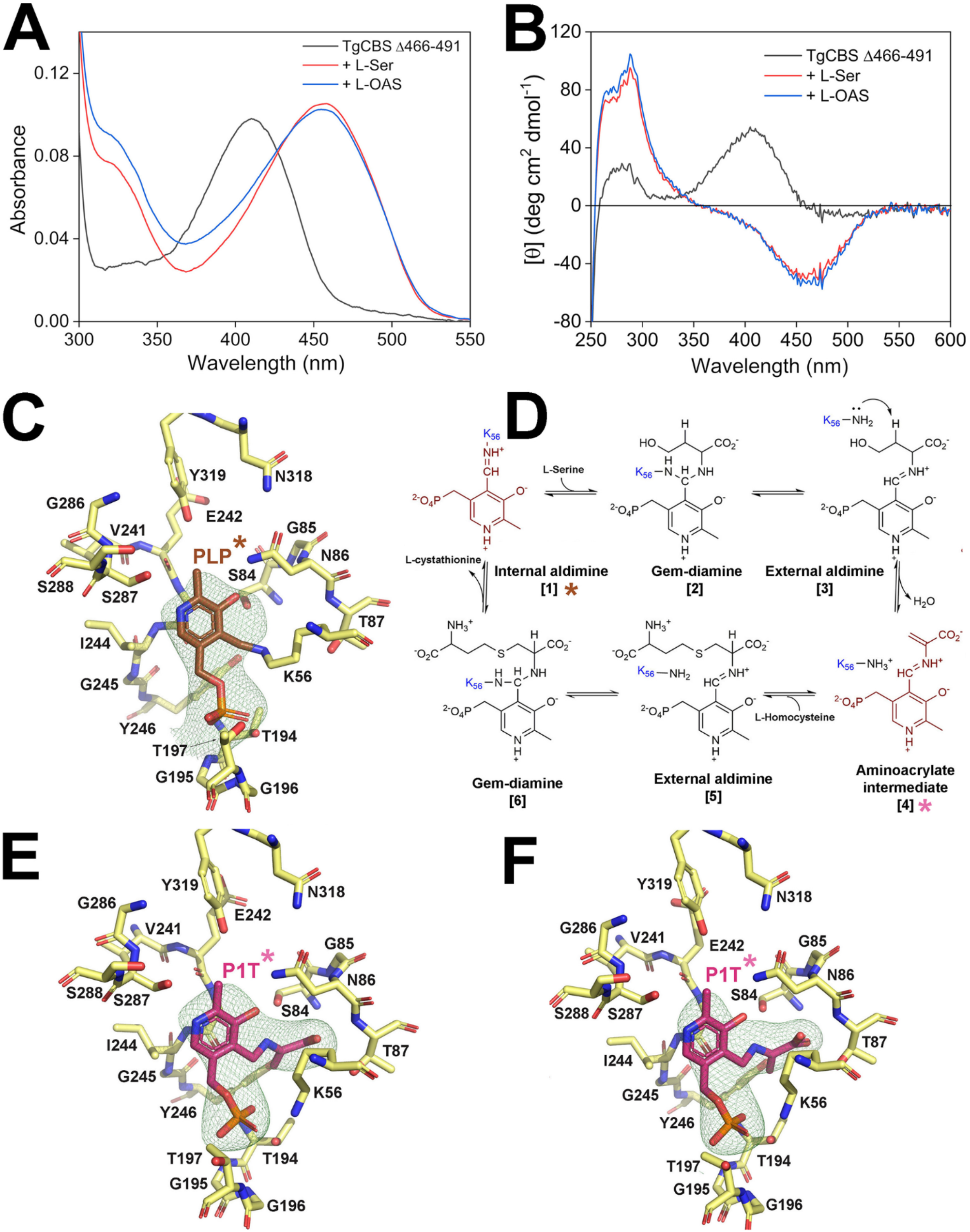
*Tg*CBS is active. (*Top*) Spectra of *Tg*CBS *Δ*466-491 with substrates. (**A**) Absorbance spectra of 15 μM *Tg*CBS *Δ*466-491 alone (*black* line) and upon addition of 10 mM Ser (*red* line) or 10 mM OAS (*blue* line). **(B)** CD spectra of 1 mg/mL *Tg*CBS *Δ*466-491 alone (*black* line) and upon addition of 10 mM Ser (*red* line) or 10 mM OAS (*blue* line). (**C**) Fo-Fc Polder omit map around PLP calculated in the absence of substrates. (**D)** Mechanism of cystathionine formation from Ser and HCys (adapted from (27)). (**C**, **E**, **F**) Zoom views of the *Tg*CBS catalytic site. The Fo-Fc Polder omit maps (contoured at 3*σ*) represent positive electron density around PLP or P1T for wt-*Tg*CBS (**C**), and 24h soaked crystals in buffer containing 10 mM Ser (**E**) or Cys (**F**). PLP and P1T are pyridoxal phosphate and aminoacrylate intermediate, respectively.

## DISCUSSION

The scarce structural information on the CBS enzyme has hindered the comparative analysis of its regulatory mechanisms across different organisms for decades. More importantly, it has significantly delayed the development of small molecules that modulate activity of this attractive therapeutic target to treat diseases such as classical homocystinuria, for which no cure is available at present. Among the five distinct domain distributions identified for the enzyme (Fig. 1C), those containing the C-terminal Bateman module are undoubtedly the ones that raise the most unknowns. Said Bateman module is present in the human enzyme, but also in that from a wide variety of pathogens, where it performs different roles that are not comprehended yet. In mammals, its role is key in regulating the activity and conformational space of the enzyme, but it seems to lack any function other than to reinforce dimeric assemblies in lower organisms such as insects and yeasts. Predicting with certainty the three-dimensional location of this intriguing domain in the overall protein fold is unfeasible at the moment. In evolutionary terms, it seemed well established that the CBS dimers can exist in two distinct conformations, mainly distinguished by the distinct orientations of their Bateman module: one basket-shaped, less active, named the *basal state* (Fig. 3B) and the other, marine mooring bollard-shaped and significantly more active, referred to as the *activated* state (Fig. 3D). The ability to adopt both patterns seemed restricted to mammals, where the transition from the first to the second necessarily requires binding of the AdoMet allosteric effector. In less evolved organisms, such as insects, only one unique constitutively activated species had been identified (27), (28) (class D in Fig. 1), (Fig. 3C), which was structurally equivalent to the AdoMet-bound CBS form of mammals (37). Intriguingly, a basket-like conformation had never been observed in any other organism until now, and its presence in *Tg*CBS was unexpected to us. Once unravelled, its structural similarity with *Hs*CBS immediately raised the question of whether *Tg*CBS exists as a unique basket-like species (Fig. 3A) or instead is capable to evolve towards a second bollard-shaped active conformation (37). We must emphasize that this knowledge is key to design efficient CBS modulators, and lays the foundation for a therapeutic intervention through the transsulfuration pathway in other organisms (c.a bacterial pathogens)(16), (17), (18). The *Tg*CBS structure unmasks for the first time the hitherto undetermined three-dimensional arrangement of class C CBSs, shared by Apicomplexa and yeast, and support that *Tg*CBS only exists as the basket-shaped species isolated in the crystals (Fig. 3A). Moreover, these findings invite to reflect on the overall three-dimensional landscape of other CBSs, as well as their structure-activity relationship.

Importantly, we have also identified the key features that constitutively retain *Tg*CBS in a basket type fold and by extension, the features that would hinder, or alternatively facilitate, the progression of other CBSs towards bollard-shaped activated states. These features reside in (i) the capacity of the Bateman module to host AdoMet at site S2 (Fig. 3) (37); (ii) the formation of interaction networks between the interdomain linker and the Bateman module that impede (or not) the displacement of the latter (Fig. 5); and (iii) the ability of the Bateman module to form an antiparallel CBS module (Supp. Fig. S3).

Based on our new data, we propose to rethink the conformational landscape of the CBSs containing a Bateman module according to three different scenarios, and not to two, as it was historically assumed so far. The first one ranges *active basal-like species* (c.a *Tg*CBS), likely not susceptible to accept and process all types of substrates and products, and without the ability to evolve conformationally to higher activity bollard-like states. In the opposite side of the conformational space are the *constitutively active and non-allosterically regulated* CBS enzymes, like those found in *D. melanogaster* (27) and *A. mellifera* (28). Finally, the third type includes the *allosterically activated* enzymes (ca. *Hs*CBS), the most evolved CBSs that that are able to adopt two different conformations, one in which the enzyme is significantly less active (*basal*) and a second one (*activated*) with high activity that can only be reached with the help of an allosteric molecule such as AdoMet, which transiently promotes the transformation (Fig. 3). Accordingly to these scenarios, enzymes with narrower access to the catalytic cavity would expectedly perform only certain reactions (either the canonical or the non canonical), limited by the entrance of fitting substrates into the catalytic cleft and/or the release of products. Of note, such tunnels access are not directly observed in the static, low-energy and highly packed crystallographic structures, but are unveiled when thermal motions and solvent effects are simulated with microsecond MD (Fig. 6), (Supp. Movies), which once again confirm the critical role of protein dynamics in enzyme catalysis (46). It remains poorly understood what type of active enzymes among the ones described above have wider catalytic ability measured under the same circumstances and by using the same methodology (an historical problem to compare the reported data on CBSs in the literature). In other words, what features award each CBS type the ability to prioritize the wide variety of reactions described so far.

Not less relevant, the structural features shared by TgCBS and the D444N CBS human variant improve our understanding of this homocystinuria causing mutant (45). Our MD analysis now reveals that subtle rotations and translations of the Bateman module are sufficient to create temporal tunnels that connect the catalytic cavity with the exterior, and that both, the size and the number of these tunnels are larger in *Tg*CBS and in the D444N variant (38) than in wild type HsCBS. The MD analysis has also revealed the dynamics of these enzymes and provides a rationale for the basal activity of wild type *Hs*CBS in the absence of AdoMet (19), which was not well explained from the proteins crystallized so far, that showed fully closed active sites, apparently inconsistent with such basal activity. Toxoplasma occupies a unique phylogenetic position since it is an early branching eukaryote, thus the study of CBS enzyme could provide useful information about the early evolution of transsulfuration routes. However, we have not found a good explanation for the ability of *Tg*CBS to process OAS in addition to Ser or Cys as it happens in related enzymes such as *Hp*OCBS (41). The comparison, both at sequence and structure level, does not show appreciable differences in the blocks of amino acids involved in the catalysis (Fig. 4), nor in the size of the catalytic cavity or the route of access for substrates until contacting the PLP cofactor. In fact, the entrance to the catalytic cavity is less compromised in the activated form of human CBS and in the insect enzymes than in *Tg*CBS due to the orientation of their respective Bateman modules. Clarifying this question will require further studies in the future.

In conclusion, our structures of TgCBS have unraveled the unique domain organization of this pivotal enzyme in the metabolism of sulfur amino acids in *Toxoplasma gondii*, the causative agent of toxoplasmosis, thus supporting the existence of an active transsulfuration pathway in the parasite. Our structures also represents the first evidence of a basket-shaped conformation in the CBS enzyme of a lower eukaryote, which breaks down barriers hitherto established about the general folding of CBS and its structure-to-function relationship across organisms. Because CBS represents a highly valuable therapeutic target, these new data paves the way for the rational design of drugs that can inhibit the activity of CBS not only in Toxoplasma but also in a wide variety of organisms, including humans. In this regard, we would like to highlight the recent discovery of the emerging role of host-generated H_2_S in bacterial pathogenesis (18), which opens the door for unprecedented host-directed H_2_S production therapies to treat microbial infections (16).

## MATERIALS AND METHODS

Unless stated otherwise, all chemicals were purchased from Sigma. *Tg*CBS construct lacking the region 466-PSTKKQAGMGEHERAKISLRKAGNSR-491 (*Tg*CBS ⊗466-491) was produced by site specific mutagenesis on the pET21a-TgCBS construct, using the QuikChange^®^ site-directed mutagenesis kit (Agilent Technologies), according to the manufacturer’s recommendations for multiple-site deletion in a single step. The primers used to introduce the deletion were as follows: forward (5’-cgtttgtgtgatggacgagaaggaatgcccacatttcctgg) and reverse (5’-ccaggaaatgtgggcattccttctcgtccatcacacaaacg). The verified plasmid was transformed into *E. coli* Rosetta (DE3) expression host cells (Novagen). The conditions for expression and purification of the mutant were as described for the wild type protein (24). The CBS activity in the canonical reactions was determined by a previously described continuous assay for Cth production (24), which employs recombinant cystathionine β-lyase (CBL) from *C. diphtheriae,* produced in our lab (47), (48) and lactate dehydrogenase (LDH) from rabbit muscle (Sigma-Aldrich) as coupling enzymes. Absorption spectra were collected on a Jasco-V560 UV-Vis spectrophotometer in 20 mM sodium phosphate pH 8.5. Far-UV CD spectra (250–200 nm) were recorded on a Jasco J-1500 spectropolarimeter as previously described (24), (40). Thermal denaturation profiles were collected by measuring CD signal at 222 nm in a temperature range from 20 to 90 °C (scan rate 1.5 °C/min). Protein concentrations was 0.2 mg/mL and measurements were performed using quartz cuvettes with a path length of 0.1 cm in 20 mM sodium phosphate pH 8.5. The oligomeric state of the *Tg*CBS construct was determined by gel filtration analysis employing a GE Healthcare Superdex 200 10/300 GL column in 20 mM sodium phosphate buffer pH 8.5, 150 mM NaCl and 0.1 mM DTT as the mobile phase as described elsewhere (24), (49). A calibration curve was constructed using the GE Healthcare high molecular weight gel filtration calibration kit, following protocols in (49).

For crystallization, the enzyme was buffer exchanged into 50 mM HEPES, 150 mM NaCl, 0.1 mM DTT pH 7.5, flash-frozen in liquid N_2_ and stored at –80°C. The crystals were grown by vapor-diffusion methods at 293 K in 96-well and 24-well crystallization plates, following a protocol described previously (50). All X-rays datasets were collected at Synchrotron beamlines XALOC (ALBA), and I03/I24 (DIAMOND, UK), and were processed with autoPROC (51). The structures were determined by molecular replacement methods with PHENIX (52) using the structure of the truncated 45-kDa *Hs*CBS (PDB ID code 1JBQ) as the initial search model. Refinement was done with PHENIX (52). Model was built with Coot (53). Figures were done with Pymol (http://www.pymol.org) and Chimera (http://www.rbvi.ucsf.edu/chimera). Sequence alignents were done with Clustal Omega (https://www.ebi.ac.uk/Tools/msa/clustalo/) and represented with Chimera. The crystal characteristics and refinement statistics are in Supp.Table S2.

### Molecular modelling and dynamics

The initial structure for the truncated human cystathionine β-synthase (D444N*Hs*CBS-Δ516-525) variant and *Tg*CBS were built by homology modelling (Schrödinger Suite) using the x-ray structure of the wild-type homodimer at 2.0 Å resolution (PDB ID 4COO for *Hs*CBS) (39) and the structure reported in this study (*Tg*CBS), respectively as templates, with the assistance of PyMol software. Residues Lys119 in both homodimer chains were covalently bound to the PLP cofactors in the form of enzyme aldimines, and parameters for these modified residues were obtained using the *antechamber* module of Amber using the *gaff2* force field and with partial charges set to fit the electrostatic potential generated with HF/6-31G(d) using the RESP (54) method. The charges were calculated according to the Merz-Singh-Kollman scheme using Gaussian 16 (http://gaussian.com/). Heme groups were modelled using all-atom parameters developed by Giammona *et. al*. (D. A. Giammona, Ph.D. thesis, University of California, Davis (1984)) Molecular dynamics (MD) simulations for each homodimer were run with the Amber 18 suite (http://ambermd.org/), using the *ff14SB* (55) and *gaff2* (56) force fields. Initial structures were neutralized with either Na^+^ or Cl^−^ ions and set at the centre of a cubic TIP3P water (57) box with a buffering distance between solute and box of 10 Å. For each complex, we followed a two-stage geometry optimization approach: the first stage minimizes only the positions of solvent molecules and ions, and the second stage is an unrestrained minimization of all the atoms in the simulation cell. The systems were then heated by incrementing the temperature from 0 to 300 K under a constant pressure of 1 atm and periodic boundary conditions. Harmonic restraints of 10 kcal mol^−1^ were applied to the solute, under the Andersen temperature coupling scheme. The time step was kept at 1 fs during the heating stages, allowing potential inhomogeneities to self-adjust. Water molecules were treated with the SHAKE algorithm (58) such that the angle between the hydrogen atoms is kept fixed through the simulations. Long-range electrostatic effects were modelled using the particle mesh Ewald method (59). A 8 Å cutoff was applied to Lennard-Jones interactions. Each system was equilibrated for 2 ns with a 2 fs time step at a constant volume and temperature of 300 K. Production was run as a 2,000 ns NVT trajectory at 300 K with a time step of 2 fs using the Andersen thermostat. Representative snapshots of the production trajectories were obtained using the *cpptraj* module of Amber and rendered with PyMol. Tunnels and channels in the protein structures were analyzed using Caver 3 software (http://www.caver.cz/) and visualized using PyMol.

## Supporting information

SuppMovieS3_TgCBS

SuppMovieS2_hCBS_D444N

SuppMovieS1_hCBS_WT

## ACKNOWLEDGEMENTS

This work was supported by Spanish Ministerio de Ciencia e Innovación (MICINN), Grants BFU2010-17857 and PID2019-109055RB-I00, Spanish Ministry of Economy and Competitiveness Grants BFU2013-47531-R and BFU2016-77408-R and BIOEF/EiTB MARATOIA BIO16/ER/035 to L. A. M.-C. IK4-TEKNIKER and CIC bioGUNE funded a PhD fellowship to CF-R, RTI2018-099592-B-C22 to GJO and PhD fellowship to RNF. We thank MINECO for the Severo Ochoa Excellence Accreditation (SEV-2016-0644) and a PhD fellowship (REF BES-2017-080435) awarded to IG-R. This study was also been supported by grants from University of Verona (FUR2018) to AA and PD. We thank the Centro Piattaforme Tecnologiche of the University of Verona for providing access to the CD spectropolarimeter.

