## Supplemental figures and tables for "Structure of Cystathionine β-Synthase from Toxoplasma gondii, a key enzyme in its H_2_S production machinery"

**Supp. Figure S1.**

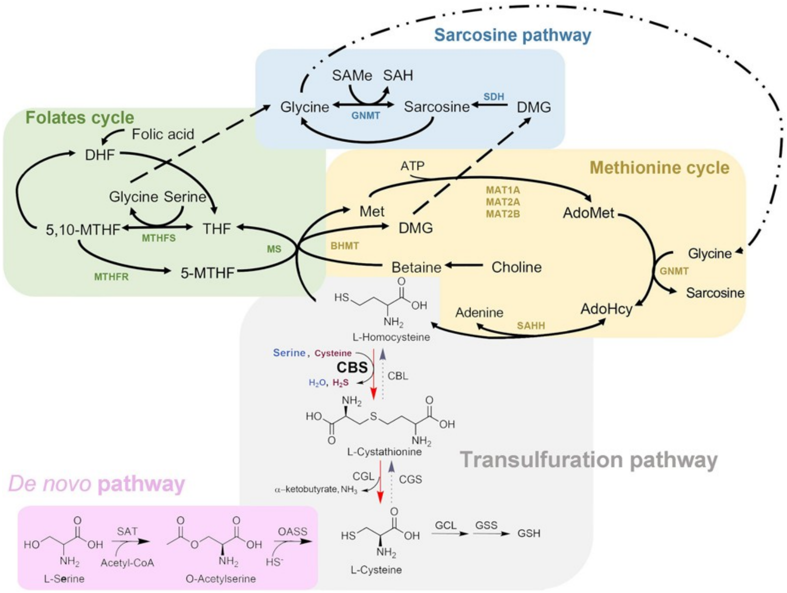

**Supp. Figure S2.**

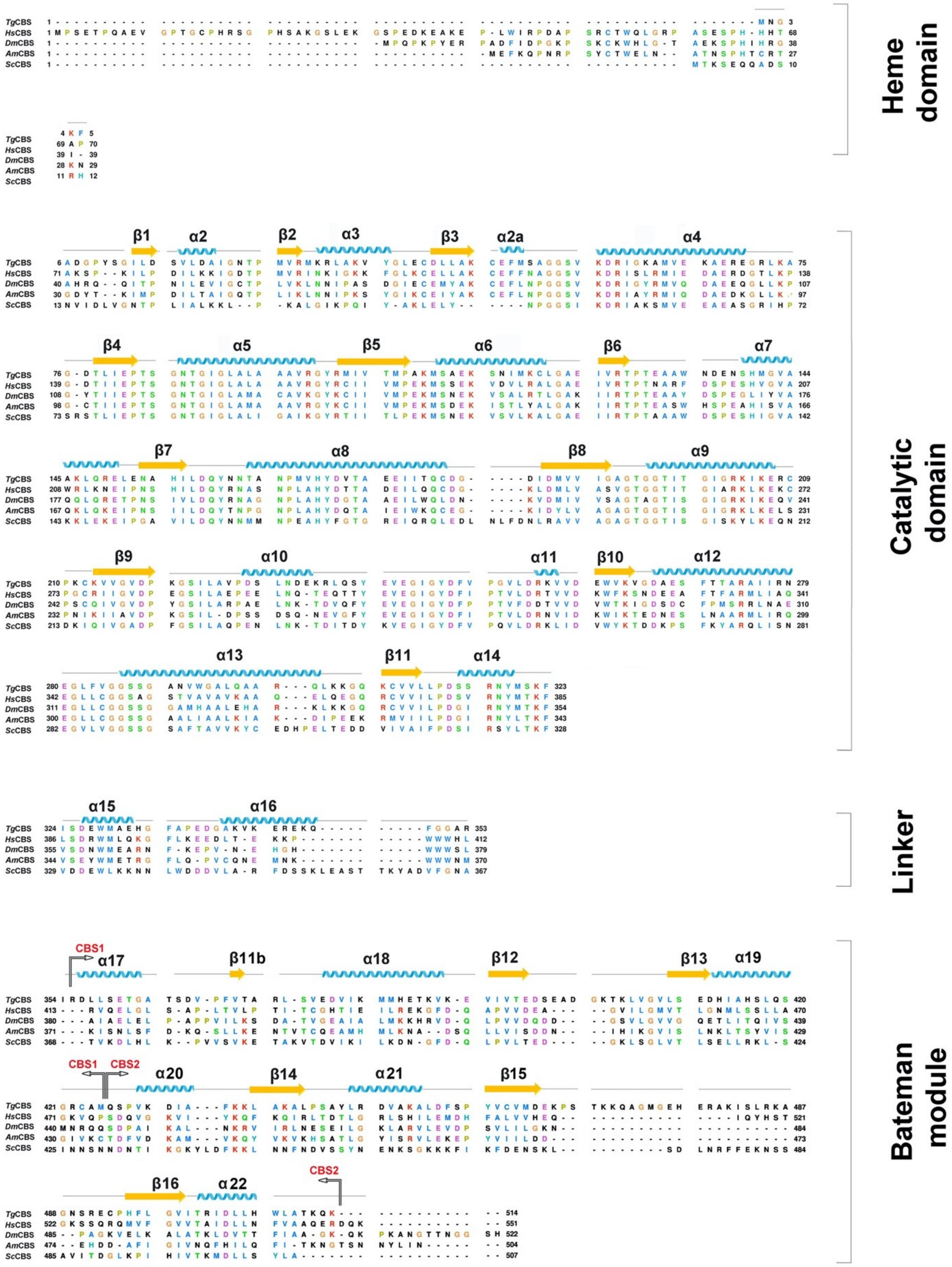

**Supp. Figure S3**

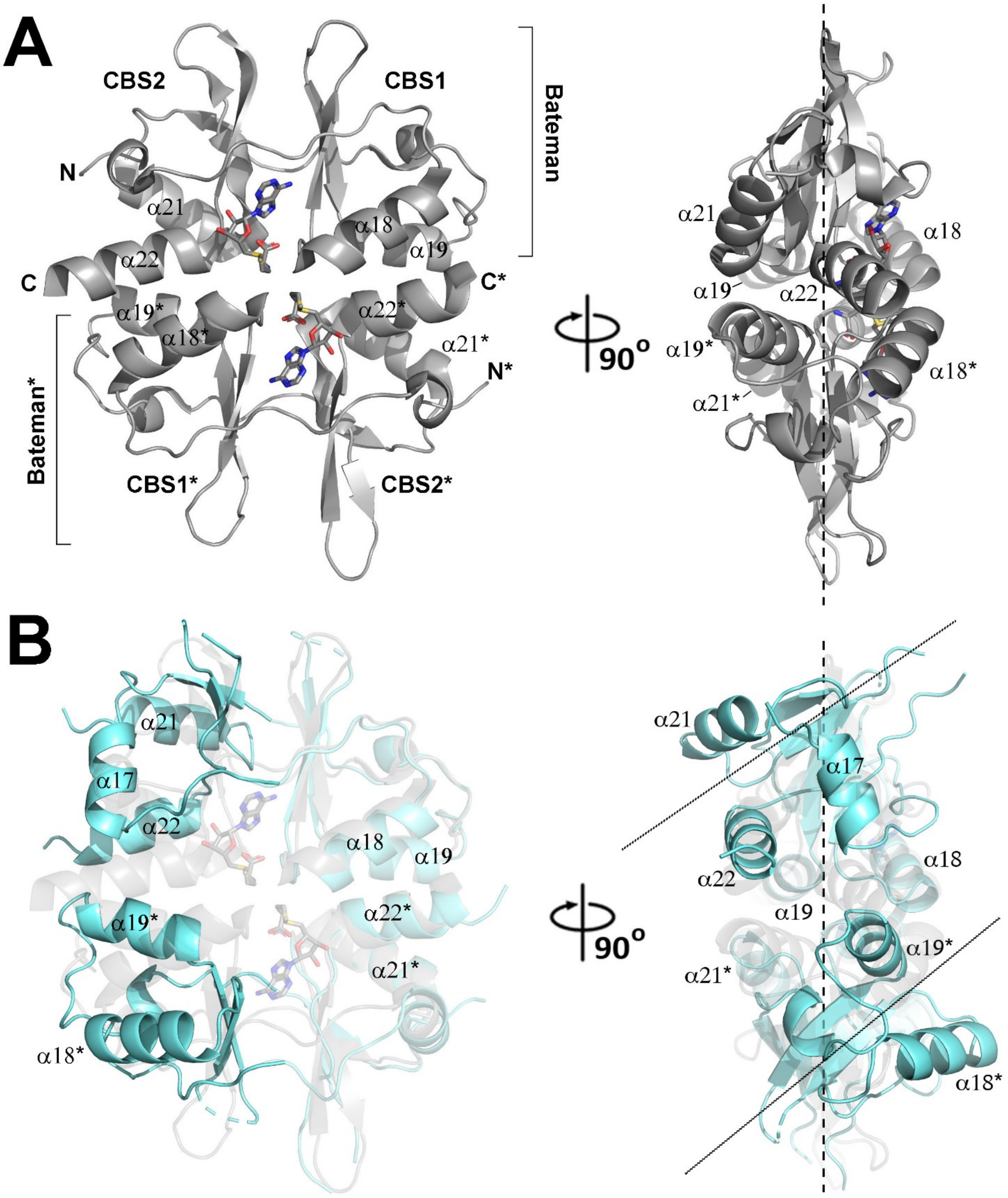

### SUPPLEMENTAL TABLES

#### Supp. Table S1.

**Steady-state kinetic parameters of TgCBS variants for canonical reactions [a].**

| Kinetic parameter | WT | Δ466-491 |
|---|---|---|
| **L-Ser + L-Hcys→L-Cth** | | |
| $k_{cat}$ ($s^{-1}$) | 6.3 ± 0.4 | 5.7 ± 0.5 |
| $K_m$ $^{L\text{-}Ser}$ (mM) | 0.42 ± 0.04 | 0.6 ± 0.1 |
| $K_m$ $^{L\text{-}Hcys}$(mM) | 0.23 ± 0.03 | 0.21 ± 0.03 |
| $k_{cat}/ K_m$ $^{L\text{-}Ser}$ ($mM^{-1}$ $s^{-1}$) | 15 ± 2 | 10 ± 2 |
| $k_{cat}/ K_m$ $^{L\text{-}Hcys}$ ($mM^{-1}$ $s^{-1}$) | 27 ± 5 | 27 ± 6 |
| $K_i$ $^{L\text{-}Hcys}$ (mM) | 1.0 ± 0.1 | 0.9 ± 0.1 |
| **L-OAS + L-Hcys→L-Cth** | | |
| $k_{cat}$ ($s^{-1}$) | 5.5 ± 0.1 | 5.6 ± 0.3 |
| $K_m$ $^{L\text{-}OAS}$ (mM) | 1.3 ± 0.2 | 1.8 ± 0.2 |
| $K_m$ $^{L\text{-}Hcys}$ (mM) | 0.20 ± 0.05 | 0.23 ± 0.05 |
| $k_{cat}/ K_m$ $^{L\text{-}OAS}$ ($mM^{-1}$ $s^{-1}$) | 4.2 ± 0.7 | 3.1 ± 0.5 |
| $k_{cat}/ K_m$ $^{L\text{-}Hcys}$ ($mM^{-1}$ $s^{-1}$) | 28 ± 7 | 24 ± 7 |
| $K_i$ $^{L\text{-}Hcys}$ (mM) | 1.4 ± 0.2 | 1.4 ± 0.3 |

**Supp. Table S2.**

| Statistics for data collection and refinement | *Tg*CBS | *Tg*CBS + Ser | *Tg*CBS + Cys |
|---|---|---|---|
| **PDB code** | 6XWL | 6XYL | 6ZS7 |
| **Space group** | $P3_1$ | $P3_1$ | $P3_1$ |
| **Unit cell** (Å)<br>a, b, c | 83.20 83.20 420.30 | 82.40 82.40 421.20 | 82.30 82.30 415.90 |
| **Resolution (Å)** | 46.10 – 3.20<br>(3.30 – 3.20) | 42.40 – 3.15<br>(3.20– 3.15) | 67.40- 3.50<br>(3.60 – 3.50) |
| **CC ½ (%)** | 99.90 (76.10) | 98.90 (68.10) | 97.70 (57.70) |
| **Redundancy** | 10.20 (9.40) | 8.30 (8.20) | 10.10 (8.00) |
| **Completeness (%)** | 79.90 (28.70) | 99.70 (99.70) | 99.70 (90.10) |
| **I/σ (I)** | 12.40 (1.99) | 7.40 (2.00) | 10.40 (2.27) |
| **Wilson B-factor (Å$^2$)** | 93.40 | 81.60 | 64.40 |
| **$R_{merge}$**[a] | 0.115 (0.999) | 0.361 (2.094) | 0.367 (1.760) |
| **$R_{meas}$**[b] | 0.121 (1.054) | 0.385 (2.395) | 0.389 (1.973) |
| **$R_{pim}$**[c] | 0.038 (0.334) | 0.133 (1.147) | 0.125 (0.624) |
| **Refinement**<br>Resolution (Å)<br>Total reflections<br>Unique reflections<br>$R_{work}$[d]/$R_{free}$[e] | <br>46.10 – 3.20<br>44707<br>42887 (2145)<br>0.253/0.277 | <br>42.40 – 3.15<br>64069<br>55325 (5571)<br>0.277/0.296 | <br>67.40 – 3.50<br>46649<br>39708 (3994)<br>0.2716/0.2818 |
| **No of non-hydrogen atoms**<br>Macromolecules<br>Ligand (PLP)<br>Ligand (P1T) | <br>21306<br>15<br>- | <br>21527<br>-<br>21 | <br>21106<br>-<br>21 |
| **Average B-factor (Å$^2$)**<br>Macromolecules<br>Ligands (PLP)<br>Ligands (P1T) | <br>89.60<br>100.40<br>- | <br>85.80<br>-<br>73.90 | <br>61.40<br>-<br>46.00 |
| **Ramachandran plot statistics (%)**<br>Res. in most favored regions<br>In additional allowed regions<br>In disallowed regions | <br>98.00<br>2.00<br>0 | <br>97.60<br>2.40<br>0 | <br>98.18<br>1.82<br>0 |
| **RMSDs**<br>Bonds length (Å)/ angle (°) | <br>0.003/0.54 | <br>0.002/0.56 | <br>0.004/0.75 |

One crystal was used for each data set. Values in parentheses are for highest-resolution shell. $R_{merge}$ [a] = $\Sigma_{hkl} \Sigma_i |I_i (hkl) - <I(hkl)>I/\Sigma_{hkl} \Sigma_i I_i(hkl)$; $R_{meas}$ [b] = $\Sigma_{hkl} \Sigma_i |I_i (hkl) - <I(hkl)>I/\Sigma_{hkl} \Sigma i\, I_i(hkl)$; [c] $R_{pim} = \Sigma_{hkl} \Sigma_i |I_i (hkl) - <I(hkl)>I/\Sigma_{hkl} \Sigma_i I_i(hkl)$; [d]$R_{work} = \Sigma |Fo - Fc| / \Sigma Fo$; [e] $R_{free} = \Sigma |Fo - Fc| / \Sigma Fo$, calculated using a random 5 % of reflections that were not included throughout refinement.

### SUPPLEMENTAL FIGURE LEGENDS

**Suppl. Figure S1. Metabolic maps.** Transsulfuration (*grey* frame) may include two opposite routes that allow the conversion of Hcys into Cys (*reverse transsulfuration, red* arrows) or viceversa (*forward transsulfuration,* dotted *grey* arrows). Transsulfuration is coupled to the folates (*green* frame) and methionine (*yellow* frame) cycles, which are coupled to the sarcosine pathway (*blue*). Some organisms (c.a. bacteria) are able to perform the *forward transsulfuration* and *de-novo* cysteine synthesis (*pink* frame). In mammals, *reverse transsulfuration* represents the sole source of Cys. Abbreviations: GSH= glutathione; OAS=*O*-acetylserine; OASS=O-Acetylserine sulfhydrylase; GCL=glutamate cysteine ligase; GSS =glutathione synthase; BHMT= betaine homocysteine methyltransferase; MS=methionine synthase; 5-MTHF= 5-methyltetrahydrofolate; THF=tetrahydrofolate; 5,10-MTHF=5,10-methylenetetrahydrofolate; MTHFS=5,10-methylenetetrahydrofolate synthetase; MTHFR=5-MTHF by methyltetrahydrofolate reductase; CGS=cystathionine $\gamma$-synthase; CBL=cystathionine $\beta$-lyase; CS=cysteine synthase; SAT=serine acetyl transferase; Met = methionine; MAT = methionine S-adenosyltransferase; AdoMet = S-adenosyl-L-methionine; GNMT= glycine N-methyltransferase; AdoHcy= S-adenosylhomocysteine; SAHH= S-adenosylhomocysteine hydrolase; CBS = cystathionine $\beta$-synthase; CGL= cystathionine $\gamma$-lyase; GCLC= glutamate-cysteine ligase; BHMT= betaine homocysteine methyltransferase; MS= methionine synthase; DHF= dihydrofolate; DMG = dimethylglycine; SDH= sarcosine dehydrogenase.

**Suppl. Figure S2. CBSs sequence alignment**. Sequence alignment of cystathionine $\beta$-synthase from *Toxoplasma Gondii* (*Tg*CBS, UniProt code A0A125YSJ9), *Homo sapiens* (*Hs*CBS, UniProt code P35520), *Drosophila melanogaster* (*Dm*CBS, UniProt code Q9VRD9), *Apis mellifera* (*Am*CBS, UniProt code Q2V0C9) and *Saccharomyces cerevisiae* (*Sc*CBS, Uniprot code P32582). The secondary elements of TgCBS are indicated.

**Suppl. Figure S3. The Bateman module of TgCBS is impaired for self-assembly**. (**A**) Disk-shaped CBS module of *Hs*CBS (PDB ID 4PCU) resulting from the assembly of complementary Bateman modules upon binding of AdoMet (in sticks). The asterisk indicates elements from the complementary subunit. The vertical dashed line represents the main plane containing each Bateman module (**B**) Superimposition of the Bateman module of *Tg*CBS (in *cyan*) on each Bateman module of the human CBS module (in *grey*). As shown, helices $\alpha$21 and $\alpha$22 do not face their equivalent complementary elements in *Tg*CBS due to a rotation of ~50 degrees of its CBS motifs with respect to *Hs*CBS. Dashed and dotted lines represent the main planes containing the CBS motifs located at the back (transparent ribbons) and the front (opaque ribbons), respectively.

**Supp. Table S1. Steady-state kinetic parameters of *Tg*CBS variants for canonical reactions [a].**[a] Reactions were carried out in 50 mM MOPS, 50 mM bicine, 50 mM proline buffer pH 9 (pH optimum) containing 0.2 mM NADH, 2 μM LDH, 1.5 μM CBL, and 0.1-30 mM mM Ser (or 0.5-100 mM OAS), 0.1-10 mM HCys and 0.2-2 μM TgCBS wild type or $\Delta$466-491 at 37°C. Data were fit as previously described (24). Reported values represent means ± S.E.M of two or more independent determinations using different batches of protein that were purified separately.

**Supp. Table S2. Statistics for data collection and refinement.**

### SUPPLEMENTAL MOVIES LEGENDS

**Suppl. Movie S1. MD analysis on wt-*Hs*CBS.** The basket-shaped basal dimer is represented in ribbons. Tunnels formed along the simulation are represented as surfaces in different colors.

**Suppl. Movie S2. MD analysis of *Hs*CBSD444N.** The basket-shaped basal dimer is represented in ribbons. Tunnels formed along the simulation are represented as surfaces in different colors.

**Suppl. Movie S3. MD analysis of *Tg*CBS.** The basket-shaped basal dimer is represented in ribbons. Tunnels formed along the simulation are represented as surfaces in different colors.
